# Independent, Ongoing Clade-Specific Expansions of IS*5* Elements in *Pseudomonas syringae*

**DOI:** 10.1101/2024.10.03.616552

**Authors:** David A. Baltrus, Audrey Sweten, Thomas Conomos, Zachary Konkel, Jonathan Jacobs

## Abstract

Insertion Sequence (IS) elements are transposable regions of DNA which are present in a majority of bacterial genomes. It has been hypothesized that differences in distributions of IS elements across bacterial strains and species may reflect underlying differences in population biology. Therefore, shifts in IS element distributions between strains may be proxies for and reflective of changes in population dynamics. Here we investigate the presence and distribution of a subclass of IS*5* elements throughout genomes of *Pseudomonas syringae*, by querying complete genomes for the presence of InsH (the main transposase found within these IS*5* elements). We report that this one subclass of IS*5* elements appears to have recently undergone multiple independent expansions in *P. syringae* clades and find that a majority of IS*5* insertion sites are not conserved across three closely related *P. syringae* pv. lachrymans genomes. We present further evidence, as has been shown for other members of the IS*5* family in different taxa, that elements from this IS*5* subclass can drive the expression of downstream genes in *P. syringae*. Taken together, our results highlight how dynamic IS*5* elements can be within and across *P. syringae* genomes and point toward the potential for IS*5* elements to rewire the expression of the *P. syringae* chromosome.

## Introduction

Insertion Sequence (IS) elements are a diverse class of transposable gene regions found throughout bacterial genomes, largely thought of as genomic parasites that proliferate within a genome [1–4]. Although IS elements are widespread throughout bacterial genomes and their importance in the creation of genetic and genomic novelty is clear, there have been relatively few investigations of their proliferation and evolutionary dynamics within and between lineages and species. Multiple factors contribute to this relative lack of studies, for instance, repetitive sequences the length of IS elements are often among the most fragmented regions when draft genomes are assembled from “short” read sequences alone and thus it can be difficult to generate accurate counts and place IS elements in proper genomic context.

Moreover, even when genomes are completely assembled, it can be challenging to compare syntenic relationships of IS elements across closely related strains. With these ideas in mind and with an interest in highlighting the evolutionary importance of IS elements, we present an analysis of the evolutionary plasticity of an IS*5* element across complete genomes of relatively closely related strains of the plant-associated and sometimes pathogenic *Pseudomonas syringae* and demonstrate the capability of this element to drive downstream gene expression in this bacterium.

IS elements are, at a minimum, composed of regions of DNA bracketed on both sides by inverted repeat sequences recognized by a transposase enzyme enabling excision/insertion of the region from/into a position in the genome [4]. In many cases, the IS element itself encodes a transposase enzyme that can recognize and excise the inverted repeat regions enabling the entire gene to autonomously “jump” or transpose to other regions of DNA within the same cell. Depending on the type of element, this jump can occur by a zero-sum “cut and paste mechanism” or can be replicative and allow for proliferation of this IS element to higher copy numbers throughout the genome. Notably, multiple IS elements can combine to form compound transposons, which can carry and transfer additional beneficial genes such as those involved in antibiotic resistance [5]. Despite numerous examples where IS element disruption is highly beneficial, the overwhelming majority of evidence suggests that IS element transpositions are predominantly either neutral or deleterious [4, 6–10]. Thus, proliferation of IS elements within and across genomes is thought to be governed by a combination of selection against transposition jumps that are deleterious coupled with genetic drift (potentially through population bottlenecks) enabling proliferation and expansion of multiple copies of these genes [11], with occasional but rare selection for strong beneficial effects.

Distributions of IS elements across genomes are hypothesized to correlate with and reflect underlying patterns in the population biology of lineages of interest. For instance, in extreme cases with populations of parasitic bacteria that are bottlenecked frequently during host transmission, the number of IS elements within a genome can explode because of excess genetic drift coupled with reduced selection pressures [12]. Lastly, elements classified in the IS*5* family are a particularly interesting group because they have been relatively well-studied for their ability to “hotwire” expression of genes and operons in proximity [13–15]. Although there have been numerous examples of transcriptional upregulation due to IS*5* elements, the specific molecular events and genomic contexts enabling them to act as promoters remain unclear [16].

*P. syringae* (sensu lato) is a bacterial species complex largely considered to be facultative phytopathogens commonly found associated with plants but whose environmental persistence and transmission may also closely be tied to the water cycle [17, 18]. There have now been thousands of genomes sequenced of strains designated within the *P. syringae* species complex. IS elements are often found within these genomes, and often disrupt genes that can contribute to virulence. However, given that repetitive regions like IS elements are also the most poorly assembled parts of the genome, there have been few large-scale assessments or general comparisons of IS element compositions across *P. syringae*. Here, we present an in-depth comparison of one subclass of IS*5* elements across complete (or nearly complete) genomes and we show that copy numbers for this particular element appear to have substantially increased independently in strains of pathovar (pv.) aesculi (a pathogen of horse chestnut) and pv. lachrymans (pathogens of cucumber) strains. While these patterns are somewhat reflective of general IS element proliferation throughout these genomes, there is no clear general signal for proliferation across all genomes. We present further evidence comparing IS*5* element positioning across three complete pv. lachrymans genomes, highlighting recent and ongoing changes in the number and position of IS*5* elements. Since IS*5* elements have the potential to act as promoters enabling transcription of downstream regions in other species, we further demonstrate the ability of this element to drive gene expression of an otherwise silent antibiotic resistance gene in one of these pv. lachrymans strains. Taken together our results suggest that multiple lineages of *P. syringae* have experienced extensive proliferation of an IS*5* element and we suggest that this genomic signature could reflect changes in population biology for this lineage compared to other *P. syringae* clades. However, it is also possible that gene expression changes and disruptions caused by this element are uniquely beneficial in the context of this pv. lachrymans and pv. aesculi pathogens.

## Materials and Methods

### IS5 Element Characterization Across Complete Genomes

The IS*5* element of *P. syringae* pv. lachrymans was originally identified from the PGAP annotation of the *Pla*107 genome. The complete protein sequence of the transposase (InsH) of this IS*5* element was used as a search query in using the DiamondBlastP function against all complete genomes at www.Pseudomonas.com with retention of hits that were >70% length of the original IS element and >98% protein identity [19]. All other comparative genomic data (specifically BlastP hits) were acquired by accessing the DiamondBlastP option at Pseudomonas.com (database version **22.1**, accessed on November 24th, 2023). Query sequences and a spreadsheet file of the BlastP results can be found at https://doi.org/10.6084/m9.figshare.27098338.v4

### Characterization of All IS Elements Across P. syringae Genomes

To further investigate IS element composition across *P. syringae* genomes, we queried all IS element classes within complete genome sequences which contained sequences for the InsH protein from blast searches as described above as well as representative complete genomes from additional phylogenetically informative strains. Insertion sequence elements were predicted using *Pseudomonas* genome assemblies via ISEscan v1.7.2.3 with default parameters [20]. Genbank accessions for each genome queried can be found as an additional file at https://doi.org/10.6084/m9.figshare.27098338.v4

### Phylogenetic Comparisons

We inferred phylogenetic relationships across all complete *P. syringae* genomes that contained an IS*5* element as well as representative complete genomes from phylogenetically informative or important clades using Realphy [21] and by designating *Pph1448a, Psy*B728a, *Pto*DC3000, *Por*1_4, *Pma*ES426 as references and with default parameters. Genome accessions for each of these assemblies and Realphy outputs can be found at https://doi.org/10.6084/m9.figshare.27098338.v4

### Comparison of IS5 Synteny and Position Across Three Complete P. syringae pv. lachrymans Genomes

For comparison across three strains of *P. syringae* pv. *lachrymans*, completely sequenced chromosomes for *P. syringae* pv. lachrymans M301350, *P. syringae* pv. lachrymans NM002, and *P. syringae* pv. lachrymans YM7902 were queried for positions of annotated InsH sequences as above using www.Pseudomonas.com. BlastP was further used to search annotated protein sequences across all three genomes using loci upstream and downstream of each identified IS*5* element sequence to identify proximate regions of synteny across the three chromosomes of these strains. If a blast search failed to retrieve hits >98% for proteins of interest (or if the query was itself an IS*5* element), we queried at least three additional loci upstream or downstream of the focal IS*5* element to establish synteny. If syntenic insertion of an IS*5* element could be clearly established by hand annotation across two or three genomes or if it was obvious that the focal copy of an IS*5* element was not present within a genome, we placed this IS*5* element into a “clear” group. If synteny could not easily be established due to rearrangements or other genomic variation and the surrounding genomic region was likely present within each genome in some context, we place the focal element into a “unclear” annotation group.

### Selection of an IS5 Element Driving Kanamycin Gene Expression

*P. amgydali* pv. lachrymans 107 (*Pla*107, also known as MAFF301315 and *Pla*N7512) was originally isolated from diseased cucumbers (*Cucumis sativus*) in Japan in 1975 and deposited at the Ministry of Agriculture, Fisheries and Forestry, Japan (MAFF no. 301315). The isolate, used to create derived strains for this report, was directly acquired from MAFF, and the complete genome assembly of this strain was originally reported in Smith et al. [22] and can be found at Genbank at accession GCA_000146005.2. Originally, a phenotypically marked version of strain *Pla*107 was created (named DAB885) through the integration of the vector pMTN1907 into a region on megaplasmid pMPPla107, and using positive selection for recombinants through tetracycline resistance [23, 24]. Although pMTN1907 also contains an *aph*3A’ locus from *Campylobacter coli* with expression driven by its native promoter, and this locus enables kanamycin resistance in *E. coli*, this gene as constituted does not enable *Pla*107 to grow on plates supplemented with kanamycin [25]. An isolate of this strain was grown overnight in KB media supplemented with tetracycline (10mg/mL) on a shaking incubator at 27°C at 220rpm. After growth overnight, 200uL of this culture was spread on KB agar plates supplemented with kanamycin (25mg/mL) and a single kanamycin-resistant colony was picked to KB liquid media. This culture was grown overnight and frozen down in the Baltrus Lab stock collection as strain DBL328. For genome sequencing, a subsample of this frozen stock was streaked to KB agar plates containing tetracycline and kanamycin, and subsequently a single colony was picked to KB liquid media supplemented with kanamycin. Genomic DNA from DBL328 was isolated from this liquid culture using a Promega (Madison, WI) Wizard kit, sequenced at Plasmidsaurus using standard protocols, and assembled using their standard pipeline involving Flye v. 2.1 [26]. The coding sequence of *aph*3A’ locus on the megaplasmid of pMPPla107 in DBL328 was queried by blastN in this assembly, with the region upstream of this locus analyzed for promoter insertions.

## Results

### Variability in Presence and Copy Number of IS5 Elements Throughout P. syringae Complete Genomes

We used DiamondBlastP to search all complete *P. syringae* (and related species) genomes through www.Pseudomonas.com and to evaluate the presence and copy numbers of annotated InsH protein (the transposase of IS*5*, encoded within this IS element). Presumably, active IS*5* elements (with intact InsH) are not universally present throughout all *P. syringae* genomes, with their presence largely localized to 3 phylogenetic clades (with stars differentiating these clades highlighted in Figure 1) from this collection of genomes in addition to the presence within a handful of scattered strains. Three of these clades are found within phylogroup 3, while one clade is found within phylogroup 1. Copy numbers within each strain are quite variable, but we note that there has been a uniform explosion to greater than 100 copies within two phylogenetic clades specifically and largely within two pathovars of *P. syringae* (pv. aesculi, a pathogen of horse chestnut; pv. lachrymans, a pathogen of cucurbits with these strains largely causative of disease in cucumber). Judging by phylogenetic relationships, and largely because of the absence of IS*5* elements within strains and clades separating *P. syringae* pv. aesculi and pv. lachrymans, a parsimonious explanation is that these IS elements independently expanded within each of these clades because there are numerous complete genome sequences that lack IS*5* elements completely interspersed in the phylogeny between these two clades.

**Figure 1.**
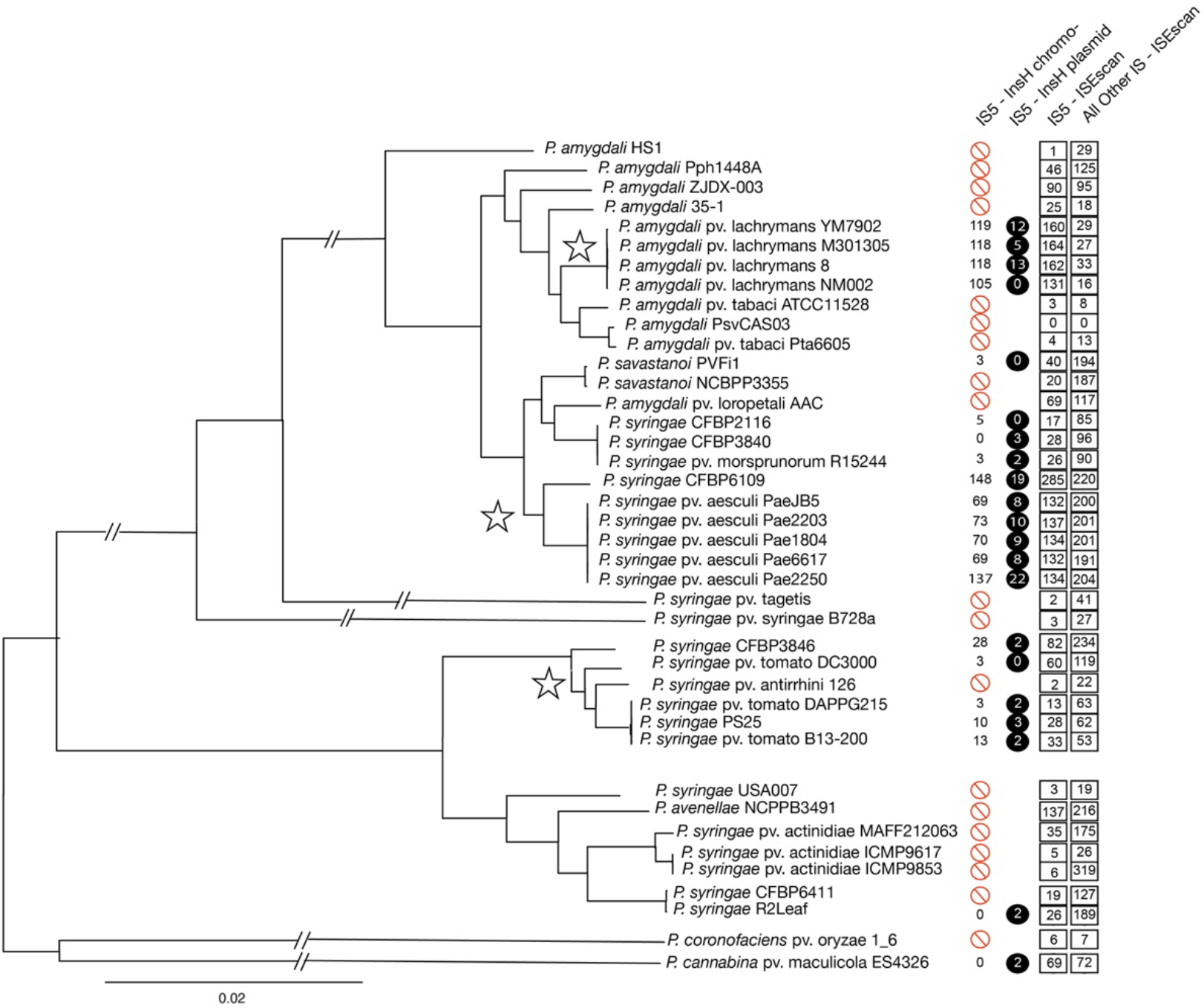
IS*5* Elements are found throughout *P. syringae* strains, with independent expansions across multiple clades. We used Realphy to infer a phylogeny across all complete *P. syringae* genomes containing the InsH transposase of the IS*5* element, while also including relevant representative complete genome sequences to adequately represent genomic diversity in this species. The first column displays either the number of IS*5* elements found on the chromosome of each strain (black numbers) or displays whether a genome lacks this element entirely (red circle). The second column (black circle with white number) displays the number of IS*5* elements that were localized to plasmids for most strains and to either plasmids or other fragments for *P. amygdali* pv. aesculi 2250. The third column displays the number of *all* IS*5* elements identified by ISEscan, while the fourth column displays the number of additional IS elements identified by ISEscan. Stars denote clades of interest for IS element expansions.

We also note that IS*5* elements are often present on the chromosome and plasmids of strains; although there are strains where the IS*5* element is present solely on the chromosome (*P. syringae* pv. lachrymans NM002, *P. savastanoi, P. syringae* pv. tomato DC3000, *P. syringae* CFBP2116) as well as strains where this element is only present on the plasmid (*P. syringae* R2Leaf, *P. syringae* CFBP3840, *P. cannabina* ES4326).

### Variability of IS Element Copy Numbers Throughout P. syringae Complete Genomes

IS element proliferation within a genome is hypothesized to potentially reflect trends in population biology for the lineages of interest. In this way, an explosion of IS*5* elements within a genome might reflect smaller effective population sizes for the lineages and clades highlighted by stars in Figure 1. Since overall population dynamics should impact copy numbers of all IS element families (rather than just IS*5*), one clear prediction is that if IS*5* element expansion is driven by changes in population biology of the strains then all IS element families should be impacted and should therefore expand in concert within these genomes. To follow up on our initial investigation of the presence of intact *insH* (and IS*5* elements) across *P. syringae*, we further queried for the presence and copy numbers of all IS element families from complete genomes containing *insH* as well as additional phylogenetically informative strains. This query also allows independent enumeration of IS*5* elements, including those that are too divergent in InsH sequence to have been included in above analyses. Although InsH protein sequences from IS*5* elements used in queries above appear to be nearly identical in sequence within and across all genomes where present, with only one or two amino acid substitutions differentiating alleles of *insH*, it is likely that there are a variety of distinct and highly diverged IS*5* elements also found within some genomes. Judging by distinct allelic classes of these IS*5* elements, it is highly likely that divergent copies have arisen from a separate horizontal gene transfer event into these strain backgrounds. From the data, there is no clear signal of correlation between IS*5* copy number and copy numbers of other IS element families within each of the lineages. Strains in *P. syringae* pv. lachrymans with large numbers of IS*5* elements contain a relatively small number of additional IS element families. Conversely, strains in *P. syringae* pv. aesculi contain large numbers of both IS*5* elements and other additional IS elements. Strains in phylogroup 1 with subtle increases in IS*5* copy numbers (between 13-30) can contain relatively few (53) to many (234) additional IS element copies. Thus, that we do not find a clear general signal that can explain IS*5* expansion. This hints that lineage level complexities in population and genome dynamics could be the main drivers of proliferation of IS elements across strains.

### Specific proliferation and movement of IS5 Elements in three P. syringae pv. lachrymans genomes

To evaluate the variability in the position of IS*5* elements across closely related strains, we compared IS*5* element distributions from the subclass queried above (represented by InsH sequences) of three *P. syringae* pv. lachrymans chromosomes. These three strains are closely related according both to phylogenetic relationships (Figure 1) and display ANI values >99% in all pairwise comparisons (see heatmap in Figure S1, calculated by FastANI at GTDB [27, 28]). Although we sought to classify conservation of each IS*5* element according to synteny across genomes, genomic variability in regions containing IS*5* elements across strains made many of these comparisons challenging. Thus, we split IS*5* elements into classes based on distributions across all three genomes and further split each locus within each class into groups (within the presence/absence split) depending on whether synteny was “clear” or “unclear”.

As shown in Figure 2, we found that 49 of the IS elements are found in the same positions across each of the three genomes, while 23 additional loci appear to be potentially conserved in position across the three strains but where syntenic relationships are unclear. Given that there are >100 IS*5* elements within each strain, a minority of all IS*5* elements in each genome are present in conserved locations across these three strains. Further, each strain possessed between 20 and30 copies of the IS*5* element which were present in unique locations across their genomes with an additional ∼10 copies per strain where it was likely that the loci were uniquely present. Many of the unique insertions were in regions of the chromosome that were variable mobile elements (predicted to be Integrated Conjugative Elements and phage regions), while numerous others were the product of local tandem duplications of a conserved IS*5* element (data not shown).

**Figure 2.**
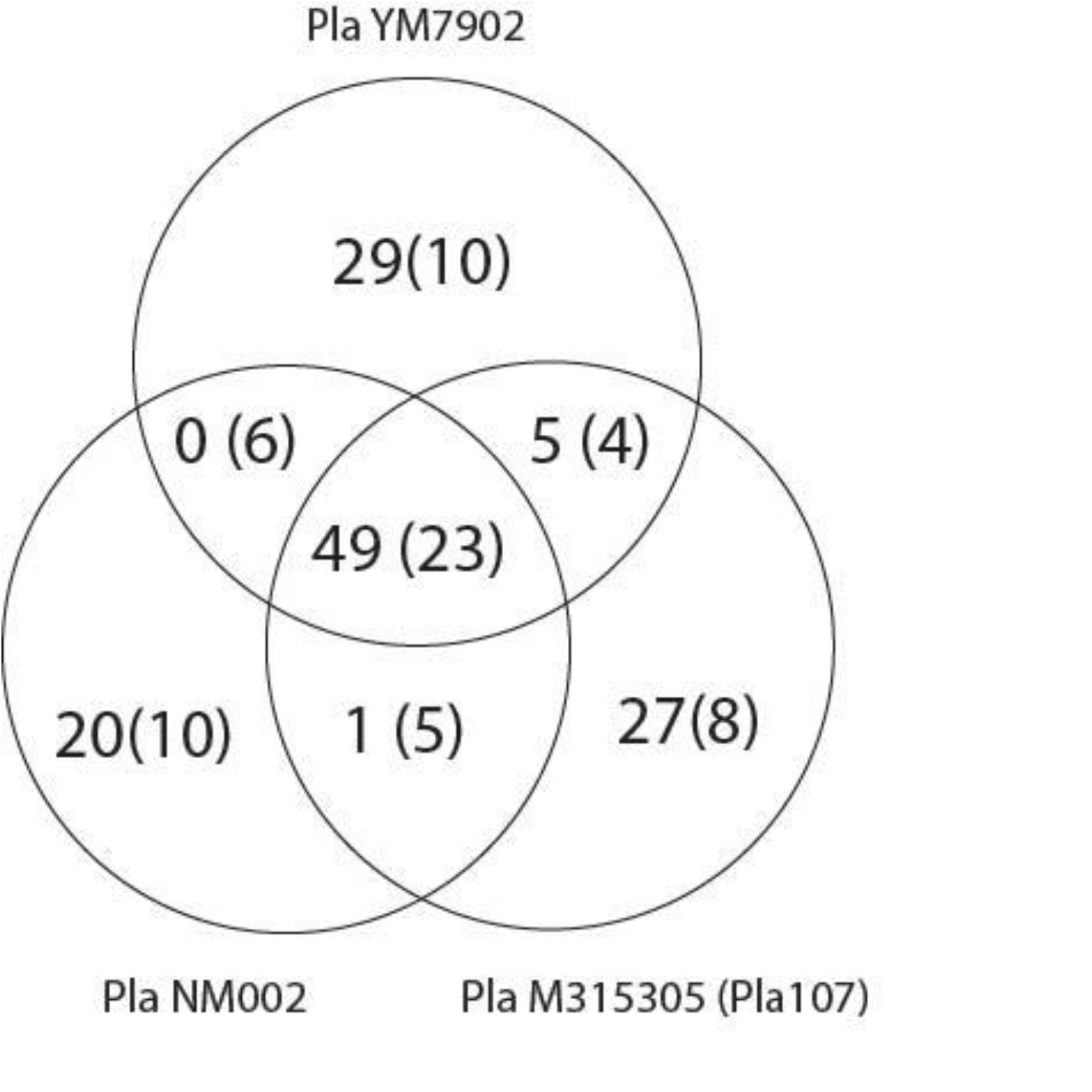
Diversity in IS*5* Element Insertion Sites Across 3 *P. amygdali* pv. lachrymans. We identified whether IS*5* element insertion points (represented by annotated InsH protein sequences) were conserved or divergent across the chromosomes of three strains within pathovar lachrymans: M315305 (*Pla*107), NM002, YM7902. The number outside of parentheses indicates situations where IS*5* elements are shared or divergent between strains with high confidence (the clear annotation group), while the number inside parentheses indicates the number of lower confidence instances (the unclear annotation group).

### *IS*5 *Elements within Pla107 are capable of driving downstream gene expression*

IS*5* elements have previously been shown to be able to drive gene expression of downstream genes in other bacterial species. We therefore took advantage of a previously constructed strain to act as a promoter trap to select this ability in IS*5* elements from *P. syringae*. The genome of *Pla*107 contains a mobilizable megaplasmid that we have previously tagged using the suicide vector pMTN1907. pMTN1907 contains open reading frames capable of encoding resistances to both kanamycin (*npt*I) and tetracycline (*tetA*), and replication of this plasmid within *Escherichia coli* provides resistance to both of these antibiotics. However, whereas the tetracycline resistance locus is expressed and provides resistance when this vector is incorporated into the genome of *Pseudomonas* strains, the *npt*I gene is not expressed in *Pseudomonas* and therefore strains with pMTN1907 recombined into their chromosomes (or other replicons) remain kanamycin sensitive. We hypothesized that differential functions of the promoter for kanamycin resistance within this plasmid could explain differential resistance phenotypes in *E. coli* and *Pseudomonas* strains. Therefore, we selected for kanamycin-resistant revertants of our previously constructed *Pla*107::pMTN1907 strain (DAB885) with the hopes that we could identify a transposition event of IS*5* elements in kanamycin resistant isolates of this strain. We found that kanamycin revertants occurred quite frequently within this strain background, and sequencing and assembly of one of these revertants demonstrated insertion of an IS*5* element upstream of the kanamycin resistance gene within the version of pMTN907 found on the megaplasmid. Specifically, it appears as though a 1210bp IS*5* family element transposed into a position 62bp upstream of the start codon for kanamycin resistance in the integrated version of pMTN1907 (Figure 3).

**Figure 3.**
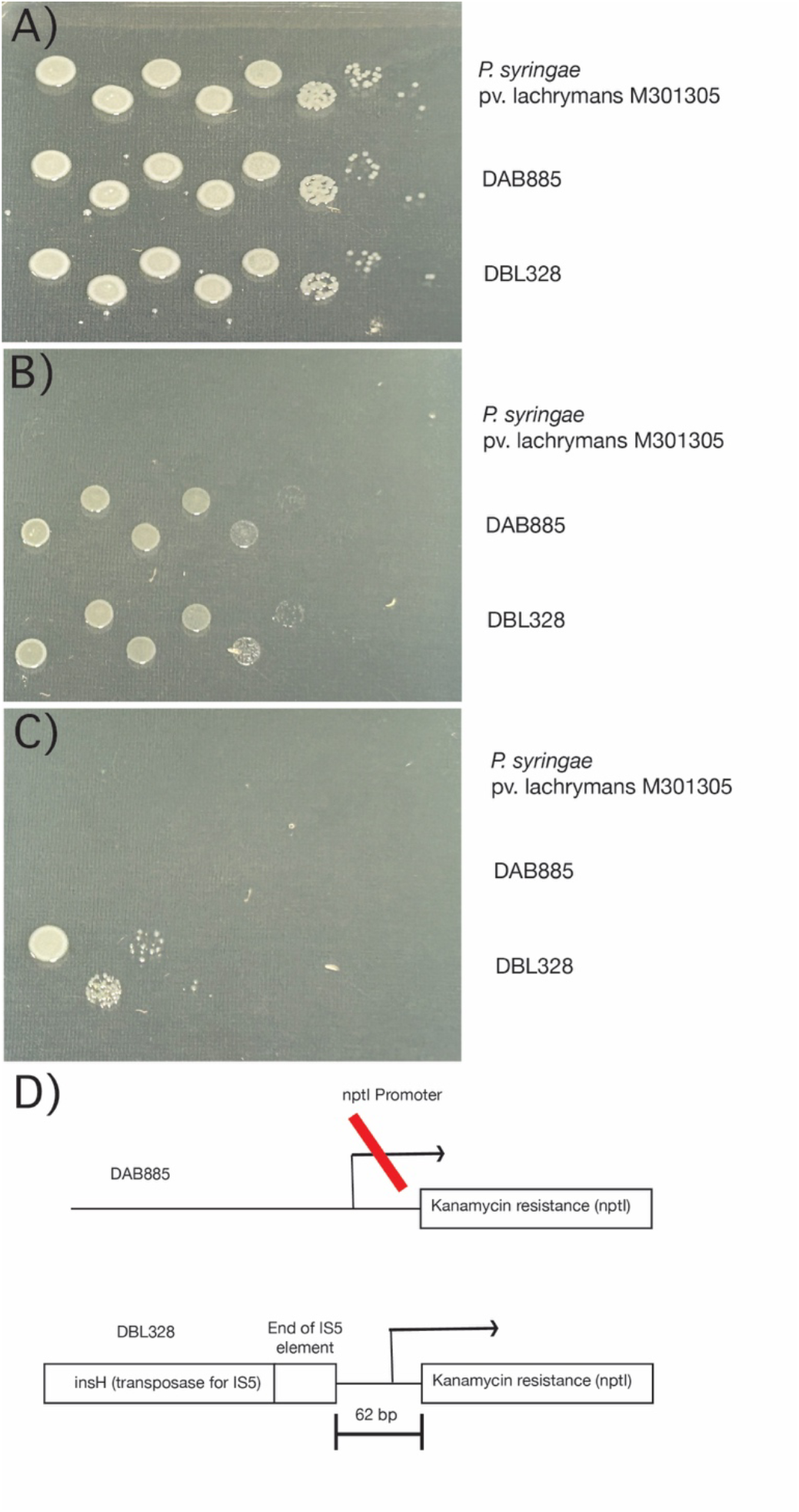
IS*5* Elements Can Drive Downstream Gene Expression. Strain DAB885 is an isolate of *P. amygdali* pv. lachrymans M301305 in which pMTN1907 has been recombined into a megaplasmid found in the strain. pMTN1907 codes for an enzyme that provides kanamycin resistance, but which is not expressed in *P. syringae*. We selected for a kanamycin resistant version of DAB885 by plating cells onto KB media containing 25ng/uL kanamycin to generate strain DBL328. A,B,C) We show growth after dilution plating overnight cultures of either *P. amygdali* pv. lachrymans M3013015, DAB885, and DBL328 onto KB media, KB media containing 10ng/uL tetracycine, or KB media containing 25ng/uL kanamycin. D) Genome sequencing demonstrated that an IS*5* element transposed upstream of the kanamycin resistance gene within DBL328 to enable kanamycin resistance.

## Discussion

### Ecological and Population Level Inferences of IS Element Proliferation

We present data demonstrating that IS*5* elements are undergoing ongoing and independent expansion across multiple phylogenetic clades of *P. syringae*. What is less clear are the evolutionary and ecological forces that enable such expansions within these specific clades. One possibility is that IS*5* elements have not expanded within other clades simply because they have been recently introduced to these clades and thus haven’t experienced a significant enough amount of time to have undergone expansion. As an argument against this relatively simple explanation, we highlight that clades which have undergone expansion (highlighted by stars in Figure 1) contain equal or less diversity across strains (are similarly aged or “younger”) than another clade with more moderate IS*5* element copy numbers.

It is also possible that there are particularities about the ecology of pv. lachrymans and pv. aesculi that enables expansion of IS elements compared to other clades. Although IS element insertions can be beneficial under certain contexts and circumstances, it is thought that these events occur in a minority of cases and that IS element copy number within a genome is driven largely by a combination of neutral and deleterious transposition events. At one end of the spectrum, transposon landing sites are restricted to regions of the genome that are not critical for carrying out cellular functions within a given environment because disruption of these regions is lethal. However, if a transposition event is non-lethal to the cell but either partially lowers fitness of the cell or is neutral, the strength of selection against this particular insertion site at a population level will be driven by population size. At relatively large effective population sizes, selection is a powerful force to cull most deleterious mutations from a population and genetic drift is comparatively weak thus limiting the fixation of neutral mutations. However, at relatively small effective population sizes, selection is much weaker and genetic drift much stronger and thus deleterious and neutral mutations can fix much more easily. Thus, it may be that pv. lachrymans strains and pv. aesculi clades possess smaller effective population sizes than other clades due to ecological differences of these strains. Perhaps they undergo smaller or more frequent population bottlenecks than other clades.

To this point, we highlight that both pv. lachrymans and pv. aesculi clades are part of phylogroup 3 of *P. syringae*. Although many *P. syringae* strains have been isolated from environmental and water sources, and it is often assumed that their life cycles are intimately tied to transmission by the water cycle, it is notable that relatively few phylogroup 3 strains have been isolated from water sources compared to other phylogroups. Thus, phylogroup 3 strains may have distinct transmission mechanisms, which could have cascading effects on their population dynamics and driving smaller effective population sizes and enabling IS5 element proliferation. While likely affected by sampling bias, most of the cases of IS element and transposon proliferation witnessed in bacterial genomes thus far are found in obligate symbionts and parasites whose population sizes are drastically bottlenecked during transmission from host to host. Flipping this idea around, differential IS element proliferation on one bacterial lineage compared to a closely related lineage could therefore be one indicator of a change of population biology in one lineage compared to another. However, we note that our data also suggests that all IS element families are not impacted similarly across each lineage. Therefore, if changes in population dynamics have led to expansions of IS*5* elements in both pvs. lachrymans and aesculi, that such expansions are not uniformly seen across all other IS elements across all lineages but only appear to have occurred in pv. aesculi strains. Thus, while data suggests that strains within pv. aesculi may display different population dynamics than other lineages of *P. syringae* as reflected in IS element copy numbers, such an explanation does not also cleanly and generally explain expansion of IS*5* elements within pv. lachrymans.

### Influence of IS Element Proliferation on Evolutionary Dynamics

Aside from possible ecological correlates of IS*5* element proliferation, what are the possible evolutionary consequences of such expansions? When IS elements proliferate throughout bacterial genomes, they seed that genome with many copies of identical ∼1000bp nucleotide sequences. These sequences can then act as landing sites for homologous recombination events, including duplications, deletions, and inversions. Therefore, regardless of changes in population dynamics that affect the fixation of mutations, *P. syringae* lineages where specific IS element families have proliferated to hundreds of copies could therefore be prone to a higher frequency of occurrence of these types of events through time and could be considered to have higher evolutionary plasticity than sister lineages without high levels of IS element proliferation. Indeed, it is worth mentioning that many of the difficulties in identifying synteny between IS*5* element insertion is that the regions of the genome that they are found in are prone to duplications, inversions, and translocations.

Additionally, one of the most intensely investigated aspects of IS*5* element biology is the ability of this family of elements to enable the expression of downstream genes and operons, and this has largely been described as occurring in multiple operons in *Escherichia coli* strains. While it is still a bit of a mystery as to how IS*5* insertion leads to upregulation, this effect appears to be more complex than simply introducing an alternative -35 or -10 hexamer upstream of the start of transcription because such effects can occur even if this element is inserted >200bp upstream of the start of transcription. Indeed, reports have implicated the global regulatory proteins H-NS and IHF as playing a role in the upregulation of operons by IS*5*. In one study, repression by H-NS was disrupted by IS*5* elements, while IHF binding correlated to bends in upstream DNA sequences promoted upregulation in another. To our knowledge, our data represents one of the clearest demonstrations that IS*5* elements can lead to upregulation in systems outside of the well-studied *E. coli* models, and could be a useful evolutionary comparison for dissecting the upregulation phenomenon at a systems level. It is currently unknown how many of the ∼>100 copy numbers of IS*5* elements actually do drive downstream gene expression in *P. syringae* pv. lachrymans and aesculi genomes but we expect future studies to better characterize the potential of these elements to change gene expression patterns.

### Conclusions

Increasing numbers of complete bacterial genomes provides an exceptional opportunity to take account of IS element variation across strains and within bacterial species and here we evaluate the distribution of IS*5* elements across the phytopathogen *Pseudomonas syringae*. Our report demonstrates that IS*5* elements are prevalent within the genomes of a variety of lineages of *Pseudomonas syringae*, that some lineages display independent expansions of this element, that these elements are actively transposing throughout these genomes such that a minority of insertion sites are conserved across closely related strains, and that this element has the potential to drive downstream gene expression within these strains. Moreover, many of the insertion sites that are unique to specific strains are present within regions of the genome which are differentially present or absent from closely related genomes, and regions of IS*5* insertions are often sites that often show inversions, and duplications/deletions. At present, it is unclear whether changes in population-level dynamics can explain the differential expansion of IS*5* elements within pvs. aesculi and lachrymans strains. It also remains unknown what fraction of the ∼100 copies of IS*5* elements within some genomes directly upregulate expression within downstream regions.

